# Functional near-infrared spectroscopy in conjunction with electroencephalography of cerebellar transcranial direct current stimulation responses in the latent neurovascular coupling space – a chronic stroke study

**DOI:** 10.1101/2020.05.24.113928

**Authors:** Zeynab Rezaee, Shashi Ranjan, Dhaval Solanki, Mahasweta Bhattacharya, MV Padma Srivastava, Uttama Lahiri, Anirban Dutta

## Abstract

Cerebellar transcranial direct current stimulation (ctDCS) can facilitate motor learning; however, ctDCS effects have not been investigated using portable neuroimaging vis-à-vis lobular electric field strength. This is important since the subject-specific residual architecture for cerebellar interconnections with the cerebral cortex, including the prefrontal cortex (PFC) and the sensorimotor cortex (SMC), can influence the ctDCS effects on the cerebral functional activation. In this study, we investigated functional near-infrared spectroscopy (fNIRS) in conjunction with electroencephalography (EEG) to measure the changes in the brain activation at the PFC and the SMC following virtual reality (VR)-based Balance Training (VBaT), before and after ctDCS treatment in 12 hemiparetic chronic stroke survivors. Furthermore, we performed general linear modeling (GLM) that can putatively associate the lobular electric field strength due to ctDCS priming with the changes in the fNIRS-EEG measures in the chronic stroke survivors. Here, fNIRS-EEG based measures were investigated in their latent space found using canonical correlation analysis (CCA) that is postulated to capture neurovascular coupling. We found that the ctDCS electrode montage, as well as the state (pre-intervention, during intervention, post-intervention), had a significant (p<0.05) effect on the changes in the canonical scores of oxy-hemoglobin (O2Hb) signal measured with fNIRS. Also, skill acquisition during first exposure to VBaT decreased the activation (canonical score of O2Hb) of PFC of the non-lesioned hemisphere in the novices at their first exposure before the ctDCS intervention. Moreover, ctDCS intervention targeting the leg representation in the cerebellum led to a decrease in the canonical scores of O2Hb at the lesioned SMC, which is postulated to be related to the cerebellar brain inhibition. Furthermore, ctDCS electrode montage, as well as the state, had a significant (p<0.05) interaction effect on the canonical scores of log10-transformed EEG bandpower. Our current study showed the feasibility of fNIRS-EEG imaging of the ctDCS responses in the latent neurovascular coupling space that can not only be used for monitoring the dynamical changes in the brain activation associated with ctDCS-facilitated VBaT, but may also be useful in subject-specific current steering for tDCS to target the cerebral fNIRS-EEG sources to reduce inter-individual variability.

## I. INTRODUCTION

Non-invasive brain stimulation (NIBS) techniques are increasingly being applied to the cerebellum for various movement disorders, including stroke [1], [2]. In these applications for stroke rehabilitation, improvement in gait and standing balance are the primary goals [3], where NIBS is usually delivered to prime the brain for learning processes by mechanisms of long-term potentiation (LTP) and long-term depression (LTD)-like plasticity [4], [5]. Cerebellar intermittent θ-burst stimulation has been shown to improve balance and gait functions in patients with hemiparesis due to stroke [6] that is important for independent gait without the risk of falling. Another NIBS technique, transcranial direct current stimulation (tDCS), is a low-cost NIBS technique amenable to home-based therapy [7] that applies a small (1-2mA) direct current (DC) via conductive electrodes placed on the scalp [8]. Recent works highlighted the adjunct role of tDCS as a modulator but not as an inducer of plasticity [9]. Due to its ease of use, especially in a home setting [10], tDCS is increasingly being investigated as an adjunct treatment to facilitate home-based stroke rehabilitation [7]. Since the cerebellum is related to movement function, especially gait and balance, as well as error-based motor learning, so cerebellar tDCS (ctDCS) has been proposed to facilitate motor adaptation during a balance learning task [11]. Zandvliet and colleagues [11] found that contra-lesional anodal ctDCS improved the standing balance performance in a tandem stance position in chronic stroke survivors but not in age-matched healthy controls. This highlighted the need for a challenging postural task to show the ctDCS effects. It was postulated that anodal ctDCS of the contra-lesional cerebellar hemisphere can strengthen the cerebellar-motor cortex (M1) connections to the affected cortical hemisphere. Zandvliet and colleagues [11] also evaluated anodal ctDCS of the ipsi-lesional cerebellar hemisphere to modulate inter-hemispheric inhibition (IHI) indirectly via cerebellar brain inhibition (CBI). However, ipsi-lesional anodal ctDCS did not improve standing balance performance in the study by Zandvliet and colleagues [11], and the neurophysiological reason remained unknown since CBI was not measured. Also, Zandvliet and colleagues [11] did not present the individual electric field distribution in the cerebellum, so the effects can be challenging to interpret without cerebellar lobule-specific dose information, especially in the elderly subjects [12]. Nevertheless, prior works on ctDCS have shown to recruit cerebellar-M1 connections depending on the ctDCS polarity [13], which can be probed with transcranial magnetic stimulation (TMS) based measure of CBI and its recruitment curve [13], [14]. However, TMS-based CBI evaluation is time-consuming and challenging to perform in a clinical rehabilitation setting where portable neuroimaging can be investigated as a proof-of-concept [15].

Cerebellar Lobules Optimal Stimulation (CLOS) pipeline [16] was developed to investigate the cerebellar lobule-specific electric field distribution for various ctDCS montages. CLOS pipeline provided age-appropriate optimization of the ctDCS electrode montage [12] for bilateral deep ctDCS of the dentate nucleus and lower-limb representations (lobules VII-IX) [17], which was proposed to facilitate standing balance in chronic stroke survivors. We first investigated the effects of ctDCS on CBI in healthy humans based on computational modeling of the electric field distribution in the cerebellum as well as neurophysiological testing [18]. We found that both anodal and cathodal ctDCS significantly decreased CBI, where the recruitment varied depending on the ctDCS polarity and the cerebellar TMS intensity used for CBI evaluation. This variability in the recruitment was postulated due to diverse effects on different cerebellar cell populations; therefore, we optimized the lobule-specific electric field distribution using CLOS for deep ctDCS of the dentate nucleus and the lower-limb representations in the cerebellum. Deep ctDCS was proposed to facilitate standing balance function during a challenging functional reach task [17] using a low-cost adaptive balance training platform [19]. Besides the cortical route, brainstem structures can also be involved since direct electrical stimulation of the cerebellum led to focal (single-joint) ipsilateral movements at the head (vermal lobule VI), face/mouth (hemispheric lobule VI), and lower-limb (hemispheric lobules VIIb-IX) [20]. The connection of the dentate nuclei (DN) with the cerebral cortex is well-known where recent work also established functional territories [21]; specifically, a functional connection was shown to the medial prefrontal cortex [21]. Importantly, viral tracing studies in non-human primates have shown Crus I and II to have projections only to the prefrontal cortex [22]. Functional neuroimaging studies, e.g., functional MRI (fMRI) [23], have shown that cerebellum projects to the association areas (prefrontal areas, posterior parietal, and superior temporal, posterior parahippocampal and cingulate areas). Recent data-driven analysis of fMRI has elucidated motor (lobules I–VI; lobule VIII), and non-motor (lobules VI-Crus I; lobules Crus II–VIIB; lobules IX–X) attentional/executive as well as default mode regions of the cerebellar function [24]. Here, prefrontal, premotor, supplementary motor, and parietal cortex have been found involved with standing balance control in hemiplegic stroke patients [25]. Although cerebellar TMS provided a method to probe the cerebellar-M1 connections in our prior work [14]; however, it was found challenging in a rehabilitation setting. Moreover, cerebellar-prefrontal connectivity cannot be probed with TMS techniques alone due to the absence of motor evoked potentials, so we investigated a portable neuroimaging approach to capture the ctDCS effects.

Based on the prior fMRI works [24], we postulated that ctDCS of the cerebellar hemispheric lobules Crus I–Crus II (in functional gradient 1, [24], [26]), the dentate nucleus, and the hemispheric lobules VIIb-IX (in functional gradient 2, [24], [26]) should modulate cerebrum activity differently during the standing balance reach task [17] that can be measured with portable neuroimaging including functional near-infrared spectroscopy (fNIRS) in conjunction with electroencephalography (EEG) [27],[28],[29]. Simultaneous fNIRS-EEG data can be used to estimate the state of the neurovascular coupling based on the tDCS-evoked responses [28], [27]. Neurovascular coupling represents interactions within the “neurovascular unit” (NVU), which consists of neuronal, glial, and vascular cells. Although focal hemodynamic effects [30] in the tDCS targeted brain areas can be due to the electric field directly affecting the neurons, astrocytes, as well as the vascular cells within the NVU [31]; yet, any remote hemodynamic effects in the brain areas unaffected by the tDCS electric field can be postulated to be driven by brain connectivity [32], [33]. Briefly, neuronal activation driven by brain connectivity will be accompanied by increased cerebral blood flow and increased cerebral metabolic rate for oxygen, leading to functional hyperemia and energy supply [16]. Vascular-based functional brain imaging techniques, such as functional MRI and fNIRS, rely on this coupling to infer changes in neural activity. The EEG can capture neural activity with excellent temporal (in milliseconds), but the poor spatial resolution (in centimeters) [35]. The fNIRS, which is less susceptible to electrical noise, is a promising tool for monitoring cerebral metabolic changes with better spatial but lower temporal resolution due to the inherent hemodynamic delay [36]. We performed fNIRS-EEG joint-imaging covering the prefrontal cortex, the primary motor cortex, and the supplementary motor area to investigate the deep ctDCS effects on the motor planning and motor control during a standing balance task. Here, it was postulated that unskilled to skilled task performance could be related to the shift in the brain activation from the prefrontal cortex (PFC) to sensorimotor cortices (SMC). On-the-fly decision-making for goal-directed behavior using our adaptive balance training platform [19] will lead to the activation of the medial prefrontal and orbitofrontal cortices during initial unskilled task performance. As the user will reach a more automatized skilled performance, the activation will shift to the sensorimotor cortices. Simultaneous fNIRS-EEG can combine the benefits from EEG and fNIRS by applying multimodal subspace signal processing to identify latent coupling variables, e.g., using canonical correlation analysis (CCA) [37], while avoiding the logistical challenges associated with fMRI-EEG in a rehabilitation setting. Here, CCA aims to maximize the correlation between EEG band power across relevant frequency band and fNIRS measure of total hemoglobin (tHb) to identify latent variables representing neurovascular coupling.

We hypothesized that a combination of fNIRS and EEG would allow for non-invasive and simultaneous assessment of neuronal activity and hemodynamic responses (i.e., neurovascular coupling response) to bilateral deep ctDCS of the dentate nucleus and lower-limb representations (lobules VII-IX). Also, we postulated that the latent variables related to the neurovascular coupling relation (found with CCA) would be differently affected at the PFC and the SMC during the adaptive balance training [19] that can be facilitated by priming the brain with bilateral ctDCS of Crus I–Crus II (in Functional Gradient 1, [24]), deep dentate nucleus and lobules VIIb-IX (in Functional Gradient 2, [24]). This hypothesis is based on prior fMRI studies [32] that showed that distinct areas of PFC are functionally connected to multiple regions of the human cerebellum, e.g., Crus I with medial prefrontal cortex (MPFC), Crus II with dorsolateral prefrontal cortex (DLPFC), while dorsal lobule VI and ventral VIIB–VIIIA with anterior prefrontal cortex (APFC). Here, Left MPFC is −12, 48, 20 and Right MPFC is 12, 48, 20; Left DLPFC is −42, 16, 36 and Right DLPFC is 42, 16, 36; and Left APFC is −32, 40, 28 and Right APFC is 32, 40, 28, in the MNI coordinate system (x, y, z). In this preliminary study, we investigated the effects of ctDCS on the relation between the latent variables (from CCA) for fNIRS and EEG at the PFC (Broadmann Areas 8, 9, and 46) using a general linear model (GLM) [38]. We also investigated the sensorimotor cortex (SMC) (Broadmann Areas 1, 2, 3, and 4) activation that is characterized by the sensorimotor rhythm, typically at a frequency of 8–13 Hz, that can be affected with ctDCS [39]. Our prior works have shown that different ctDCS electrode montages affect different parts of the cerebellum [7,21, including effects on the standing balance performance in chronic stroke survivors using a Generalized Linear Model [17]; however portable neuroimaging of the ctDCS response was not presented vis-à-vis lobule-specific electric field. In this proof-of-concept study, we evaluated the feasibility of fNIRS-EEG joint-imaging in the chronic stroke survivors (without cerebellar lesion) at the point of care in low-resource settings [17].

## II. Methods

### A. Subjects and their age-appropriate head modeling for deep ctDCS

A convenience sample of a total of 14 male chronic (>6 months’ post-stroke) stroke subjects volunteered for the study. The hemiplegic subjects, who (i) were aged between 18 and 90 years, (ii) could walk independently for at least 10 meters, (iii) could provide informed, and written consent, and (iv) could understand instruction from the experimenter, were enrolled. We then selected chronic stroke survivors with cerebral lesions but with an intact cerebellum so that the ctDCS electric field effects can be delivered to the cerebrum via the cerebellum [42]. Out of these 14 male volunteers, one (P13) could not complete the study because of his unavailability after initial screening, and the other (P14) was excluded from the study based on the exclusion criteria. Table 1 lists the 12 male chronic stroke subjects (P1-P12, Mean (SD) = 46(±13) years) who participated in the current study. T1-weighted MRI was available only for the subjects P1-P6 (from All India Institute of Medical Sciences New Delhi, India) (details provided in Zeynab et al. [16]). Written informed consent was obtained from each subject, and the multi-center research protocol for this study was approved by the All India Institute of Medical Sciences, New Delhi, India Institutional Review Board (IEC-129/07.04.2017), and Indian Institute of Technology Gandhinagar, India Institutional Review Board (IEC/2019-20/4/UL/046).

**Table 1:**
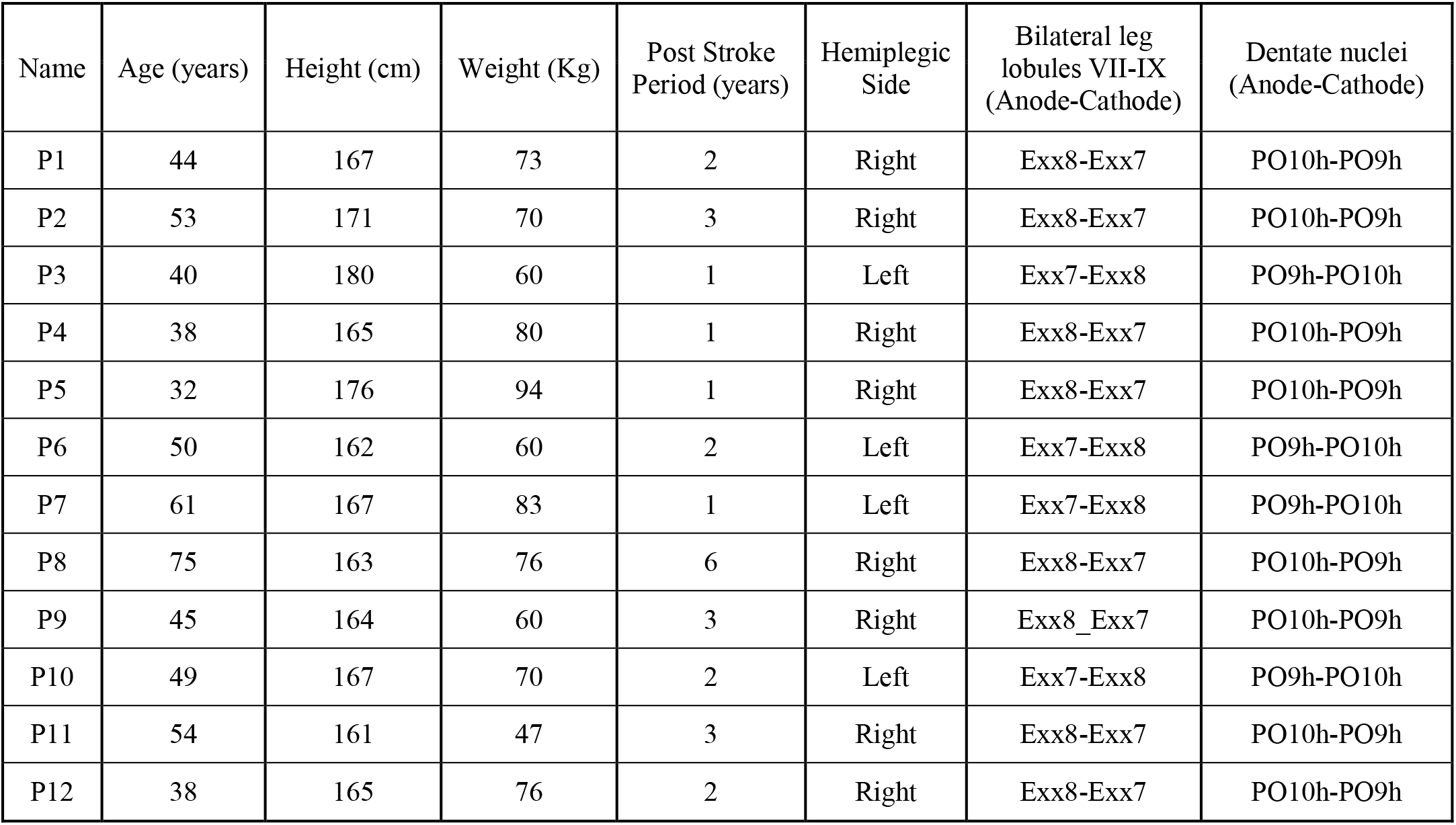
Subject characteristics (all male) and subject-specific bipolar ctDCS

### B. Computational head modeling and flatmap visualization

Computational head modeling (details provided in Zeynab et al. [16]) was performed using age-appropriate averaged (n = 73) human brain MRI template for the relevant age-groups (see Table 1) that were obtained online at the https://jerlab.sc.edu/projects/neurodevelopmental-mri-database/ with the permission of Dr. John Richards. The subjects for the human brain MRI templates were all normal healthy adults with no history of neurological or psychiatric illness, head trauma with loss of consciousness, or current or past use of psycho-stimulant medications, cardiovascular disease, and no abnormal findings on the MRI [43]. Neurodevelopmental MRI Database [43–47] consisted of average (male and female) T1-weighted MRI for the head across different age-groups as well as the segmentation priors for gray matter (GM), white matter (WM), and cerebrospinal fluid (CSF), which was used for head modeling in CLOS pipeline [16]. The selected age-groups are as follows: (N is the number of subjects’ MRI, which were used to average for that age-group), P1: 40-44 years (N=61); P2: 50-54 years (N=57); P3: 40-44 years (N=61); P4: 35-39 years (N=50); P5: 30-34 years (N=63); P6: 50-54 years (N=57); P7: 60-64 years (N=83); P8: 75-79 years (N=61); P9: 45-49 years (N=65); P10: 45-49 years (N=65); P11: 50-54 years (N=57); P12: 35-39 years (N=50). Tetrahedral volume mesh was created using the ROAST toolbox [48], which is a Matlab (Mathworks, MA, USA) script based on three open-source software; Statistical Parametric Mapping (SPM) [49], Iso2mesh [50], and getDP [51]. ROAST used SPM12 [52] to segment the head and the brain. After SPM12 segmentation and manual comparison with the segmentation priors from the Neurodevelopmental MRI Database [43–47], five tissues were labeled for the tetrahedral volume mesh, namely, Scalp, Skull, CSF, GM, and WM. These different brain tissues for the volume mesh were modeled as different volume conductors during finite element analysis (FEA) in the ROAST. Here, isotropic conductivity based on prior works was used for different brain tissues [53], which were (in S/m): Scalp = 0.465; Skull = 0.01; CSF = 1.654; GM = 0.276; WM = 0.126 [7,41–43]. In all the simulations, the voxel size was 1mm^3^. Computational modeling of the cerebellar lobule-specific electric field distribution was performed for two bipolar ctDCS montages (details provided in Zeynab et al. [16]). To investigate the electric field distribution in the cerebellar lobules, we extracted the electric field distribution within cerebellar white and gray matter. The electric field was extracted using the CLOS pipeline and cerebellar masks of each head model assessed from the SUIT toolbox. We defined parcellation of the cerebellar regions according to the SUIT atlas. The cerebellar maps were visually inspected in an image viewer software. The 2D representation of electric field distribution and visualization of FEA data across cerebellar volume for all patients was implemented by using the flatmap script in the SUIT toolbox.

#### (1) Bipolar PO9h – PO10h montage for dentate nuclei

A 3.14cm^2^ (1cm radius) circular anode was placed at the contra-lesional side, and a 3.14cm^2^ cathode was placed at the ipsi-lesional side for ctDCS with 2mA direct current.

#### (2) Bipolar Exx7 – Exx8 montage for bilateral leg lobules VII-IX

A 3.14cm^2^ (1cm radius) circular anode was placed at the contra-lesional side, and a 3.14cm^2^ cathode was placed at the ipsi-lesional side for ctDCS with 2mA direct current.

### C. Experimental protocol for functional near-infrared spectroscopy and electroencephalography (fNIRS-EEG) of ctDCS effects in the latent canonical space

At the start of the study session, the subjects were asked to sit and relax for about 5 minutes. Then, the experimenter explained to the subject what he was expected to do in the study. The study required a commitment of about an hour. In a counterbalanced cross-over repeated measure experimental design, bipolar ctDCS was performed with montages, Bipolar PO9h – PO10h, and Bipolar Exx7 – Exx8 (see section B for details). The experimenter prepared the subject by placing an fNIRS-EEG/tDCS STARSTIM 8 (Neuroelectrics, Spain) cap on the subject’s head. Then, 2 minutes of baseline fNIRS-EEG data were collected in the resting-state. Then, the subjects performed virtual reality (VR)-based Balance Training (VBaT) [19] for about 10 minutes. During VBaT, subjects had to shift the body weight in different directions (North, South, East, West, etc.) using an ankle strategy based on the cues from the VBaT platform [19]. After this, the subject was asked to sit and relax on a chair when 2 mins of fNIRS-EEG data was collected. Subsequently, bipolar ctDCS was delivered for 15 minutes in the seated resting position with a current intensity of 2 mA via 3.14 cm^2^ Pistim (Neuroelectrics, Spain) electrodes that were also used for EEG monitoring. Following this, 2 mins of post-tDCS fNIRS-EEG data were collected before the subject repeated VBaT tasks, followed by another 2 mins of fNIRS-EEG recording in the resting-state. Therefore, the fNIRS-EEG data [56] were recorded in continuous epochs of 30 seconds for 2 minutes before and after VBaT tasks, which were conducted before (pre-intervention) and after (post-intervention) 15 minutes of bipolar ctDCS. Figure 1 shows the experimental setup and the protocol.

**Figure 1:**
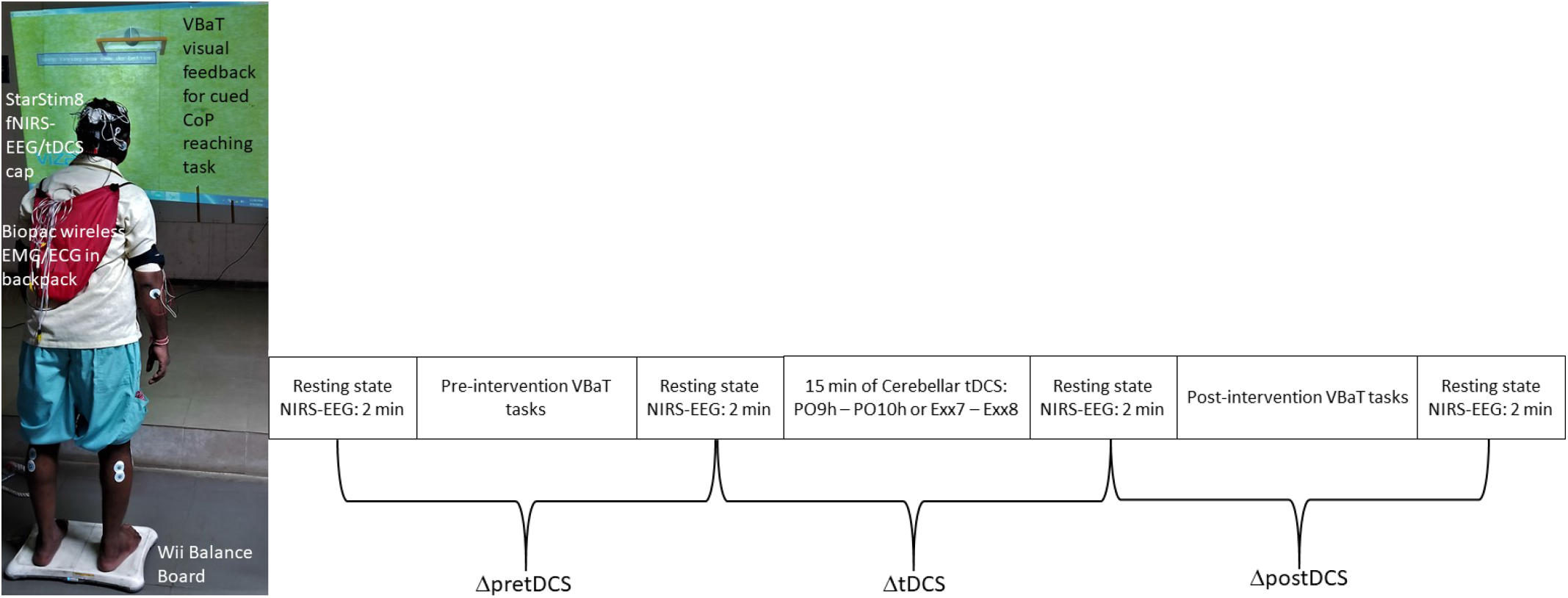
Left panel: Experimental setup for ctDCS facilitated virtual reality (VR)-based Balance Training (VBaT). Right panel: experimental protocol - resting-state fNIRS-EEG data was collected for 2 minutes before and after the measures of virtual reality-based Balance Training (VBaT) before (pre-intervention) as well as after (post-intervention) 15 minutes of cerebellar anodal tDCS using two bipolar montages (PO9h – PO10h or Exx7 – Exx8) on 12 stroke survivors using a counterbalanced cross-over repeated measures experimental design

The fNIRS-EEG portable neuroimaging and triaxial accelerometry (at 100 S/s) were conducted using a wireless fNIRS-EEG system [31] that used the lab streaming layer (LSL) for time-synchronization. STARSTIM fNIRS-EEG system combined a wireless STARSTIM 8 stimulator (Neuroelectrics, Spain) for 8-channel tDCS and EEG in conjunction with fNIRS with the Octamon+ system (Artinis, Netherlands). The 8-channel EEG was recorded from Fp1, Fp2, F3, F4, C3, C4, P3, P4 scalp locations (10–20 positioning system). The eight fNIRS sources were positioned at AF7, AF3, AF8, AF4, CP4, FC4, CP3, FC3, and the two fNIRS detectors were placed at the Cz and FPz with a source-detector distance of around 35mm. Computational head modeling for fNIRS sensitivity analysis was also performed with an age-appropriate averaged human brain MRI template. Sensitivity analysis was performed for the fNIRS montage consisting of 8 sources and two detectors using Monte Carlo simulation that provided a probabilistic model of the path of the photons. Monte Carlo simulation generated a forward matrix that represented the spatial sensitivity profile of each channel (source-detector pair) to the cortical absorption changes. Here, the sensitivity analysis was performed using the AtlasViewer toolbox [58], which is a toolbox for the determination of brain coordinates along with the sensitivity measurement for the source-detector montages. AtlasViewer toolbox facilitates optimal placement of the optodes or probes for the fNIRS studies. The probes were first designed in AtlasViewer with the eight sources placed at AF7, AF3, AF8, AF4, CP4, FC4, CP3 FC3, and the two detectors placed at Cz and FPz. The probe design was then registered to the surface of the individual head model, followed by the projection to the cortex. This projection to the cortex, found by drawing vectors perpendicular to the surface at the midpoint of each source-detector pair, ensured the sensitivity to the brain regions including PFC (Broadmann Areas 8, 9, and 46), and SMC (Broadmann Areas 1, 2, 3, and 4). The Monte-Carlo simulation was set up with 100 million photons being injected by each source using the default optical parameters for each layer of the head model. The sensitivity profile is shown in Figure S1 (in Appendix) where the voxel size was considered as 1mm^3^.

The VBaT tasks required the subjects to maneuver different virtual objects in the VR environment using their center of pressure (CoP) while performing weight shifting on a Wii balance board (WiiBB) and then maintaining their posture to hold the virtual objects at target locations in the VR space [19]. The CoP values were acquired from WiiBB at 30 Hz and were preprocessed by a 5-point moving average filter before being used to modulate the virtual objects in the VR environment. Successfully reaching the target using CoP provided audio-visual feedback to the subjects. The resting-state fNIRS-EEG data before and after the VBaT were preprocessed with the HOMER3 (https://github.com/BUNPC/Homer3) and the EEGLAB toolbox [32] in Matlab (Mathworks, Inc. USA). EEG data was first down-sampled to 250Hz and then was de-trended by high-pass filtering with 1 Hz cutoff frequency, line noise, and its harmonics were removed using “CleanLine,” and then EEG data were re-referenced to the common average. Then, the continuous EEG data were processed with Artifact Subspace Reconstruction (ASR) [60], and then Adaptive Mixture Independent Component Analysis (AMICA) was performed in EEGlab [61]. The artifactual components were removed by IClabel [62] and visual inspection [63], which made the fNIRS-EEG data discontinuous with the discontinuity marked as events in the EEGlab. We used HOMER3 routines in Matlab (Mathworks, Inc.) for the fNIRS data analysis based on the modified Beer-Lambert law to find the changes in the oxy-hemoglobin (O2Hb) and deoxy-hemoglobin (HHb) concentrations. One of the challenges is a high sensitivity for the scalp and other extracranial hemodynamics, including changes in the heartbeat and the breathing cycle, that can mask cerebral activation [64]. Usually, more significant changes are observed in the extracranial O2Hb when compared to extracranial HHb concentration. First, channels with low scalp coupling were identified based on PHOEBE approach [65] that flagged most of the VBaT data. Although one can apply HOMER3 method (https://github.com/avolu/tCCA-GLM) proposed by Luhmann et al. [66] using synchronized triaxial accelerometer data (from STARSTIM 8) to reduce the movement artifacts; here, we discarded all the VBaT fNIRS-EEG data due to possible detrimental effects of co-modulated noise on the NVC analysis (CCA). Thereafter, we performed band-pass (0.07Hz-0.13Hz) Butterworth filtering to remove the heart rate, respiratory rate, and other high and low-frequency noise. Subsequently, we performed principal component analysis (PCA) to remove the global covariance due to any global interference by the extracranial hemodynamic signals [42]. After preprocessing the fNIRS-EEG data in the EEGlab and HOMER3, we identified four epochs of continuous 30 seconds of resting-state fNIRS-EEG data (using the discontinuity event markers) before and after VBaT tasks.

A total of 4 groups of 2 minutes of fNIRS-EEG data were recorded in the resting state conditions before and after VBaT task, which were performed before and after the bipolar ctDCS intervention (see Figure 1 for the experimental protocol). Therefore, fNIRS-EEG signal changes (*i. e.*, 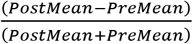) could be measured from before to after VBaT task before ctDCS intervention (called (Δ*pretDCS*), from before to after ctDCS intervention (called Δ*tDCS*), and from before to after VBaT task after ctDCS intervention (called Δ*postDCS*). We evaluated NVC between the log10-transformed EEG bandpower within 1-45Hz, and the total hemoglobin (tHb=O2Hb+HHb) signals within the 0.07Hz-0.13Hz based on canonical correlation, as described next. Temporally embedded Canonical Correlation Analysis (tCCA) was applied separately to the fNIRS-EEG data from the left and the right prefrontal cortex (PFC) (fNIRS – left: AF3-FPz, AF7-FPz; right: AF4-FPz, AF8-FPz, and EEG – left: Fp1, F3; right Fp2, F4) as well as the left and the right sensorimotor cortex (SMC) (fNIRS – left: CP3-Cz, FC3-Cz; right: CP4-Cz, FC4-Cz, and EEG left: C3, P3; right C4, P4). Here, tCCA maximized the correlation between the log10-transformed EEG bandpower and fNIRS O2Hb, HHb, and tHb (=O2Hb+HHb) to identify the latent variables (or, canonical scores) where the canonical correlation between the log10-transformed EEG bandpower, and the tHb (blood volume) was used as a measure of NVC. Also, the canonical scores for fNIRS O2Hb and HHb signal, as well as the log10-transformed EEG bandpower, were used as an indicator of the change in the brain activation at the PFC and SMC locations. Here, the canonical scores for fNIRS O2Hb and HHb signal in the latent space are postulated to be less sensitive to the systemic artifacts than the fNIRS O2Hb and HHb signal themselves. We performed an analysis of variance (ANOVA) to test the effects of the factors, brain location, ctDCS montage, and state (Δ*pretDCS*, Δ*tDCS*, Δ*postDCS*), on the changes (*i. e.*, 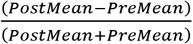) in the average NVC and canonical scores using a cross-over repeated measure design at a 5% significance level.

### D. Multivariate general linear model with lobular electric field distribution as the predictor, and the brain activation as the response variables

In a counterbalanced repeated-measure cross-over design to investigate the ctDCS effects, we investigated the effects of the lobular electric field distribution in the cerebellum on the changes in the brain activation (*i. e.*, 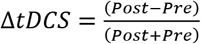) using a multivariate general linear model (GLM). The following changes in the brain activations were measured in the resting state at the PFC and SMC locations of the ipsi-lesional (Ipsi) and contra-lesional (Contra) hemispheres: canonical scores for fNIRS O2Hb, HHb, and log10-transformed EEG bandpower signal. The average strength of the electric field in the following cerebellar locations were based on computational modeling using ipsi-lesional (Ipsi) and contra-lesional (Contra) SUIT labels [16]: ‘Ipsi I-IV’, ‘Contra I-IV’, ‘Ipsi V’,’Contra V’, ‘Ipsi VI’, ‘Vermis VI’, ‘Contra VI’, ‘Ipsi Crus I’, ‘Vermis Crus I’, ‘Contra Crus I’, ‘Ipsi Crus II’, ‘Vermis Crus II’, ‘Contra Crus II’, ‘Ipsi VIIb’, ‘Vermis VIIb’, ‘Contra VIIb’, ‘Ipsi VIIIa’, ‘Vermis VIIIa’, ‘Contra VIIIa’, ‘Ipsi VIIIb’, ‘Vermis VIIIb’, ‘Contra VIIIb’, ‘Ipsi IX’, ‘Vermis IX’, ‘Contra IX’, ‘Ipsi X’, ‘Vermis X’, ‘Contra X’, ‘Ipsi Dentate’, ‘Contra Dentate’, ‘Ipsi Interposed’, ‘Contra Interposed’, ‘Ipsi Fastigial’, ‘Contra Fastigial’, ‘Non-Cerebellar Brain’. Due to multicollinearity and singularity across the lobular average electric field strengths, principal component analysis (PCA) was conducted to identify principal components (PCs) that accounted for greater than 98% variance across subjects. For principal component regression, a multivariate GLM (’mvregress’ in Matlab) was fitted to the PCs of the mean lobular electric field strength as the predictor and the latent variables of the brain activation as the response.

## III. Results

### A. Computational head modeling

We investigated the 3D lobular electric field distribution using 2D visualization of the simulation results in SUIT flatmap – see Figure 2a. The SUIT flatmap for the 3D lobular electric field distribution in the cerebellar lobules is shown in Figure 2b for the twelve subjects. The results for the bipolar PO9h – PO10h montage for dentate nuclei is shown in the right column of Figure 2b, and the result for the bipolar Exx7 – Exx8 montage for bilateral leg lobules VII-IX is shown in the left column of Figure 2b.

**Figure 2 :**
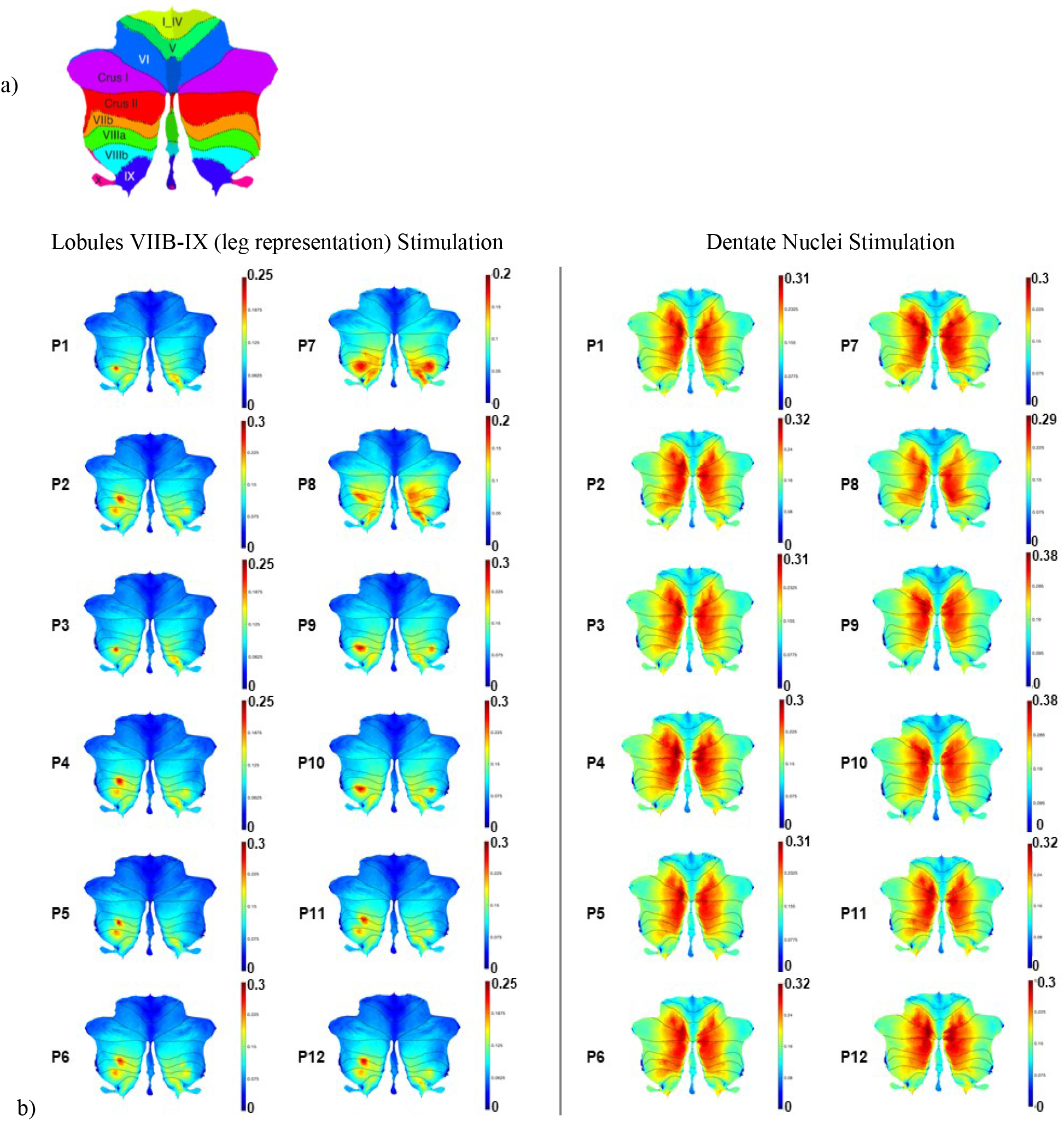
a) 2D representation of the 28 cerebellar lobules using SUIT flatmap. b) Electric field strength (V/m) across different subjects using SUIT flatmap (SUIT toolbox) with the color scale representing the electric field strength shown on the right. Left two columns show the electric field distribution that resulted from the bipolar Exx7 – Exx8 montage for bilateral leg lobules VII-IX while the right two columns present the results for the bipolar PO9h – PO10h montage for dentate nuclei.

### B. Functional near-infrared spectroscopy and electroencephalography in the latent (canonical) space

Three-way ANOVA test on the effects of the factors, brain location, ctDCS montage, and state (Δ*pretDCS*, Δ*tDCS*, Δ*postDCS*), showed a moderate interaction effect between ctDCS montage and state on the canonical scores of O2Hb (p=0.0788). Also, a significant effect of the state (p=0.0019) as well as the interaction between ctDCS montage and state (p=0.0153) was found on the canonical scores of log10-transformed EEG bandpower, as shown in Figure 3.

**Figure 3:**
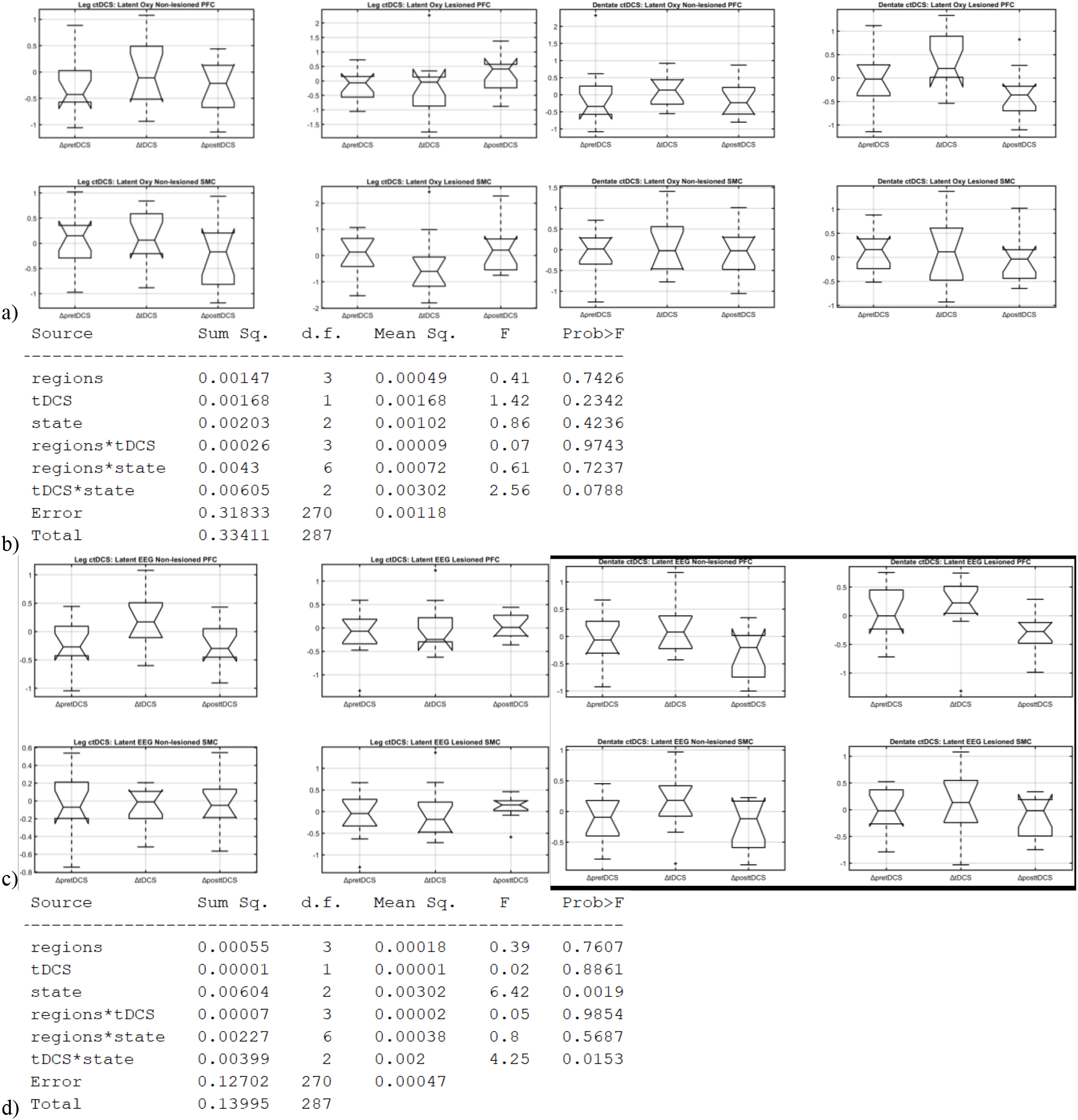
Three-way ANOVA results shown along with the boxplots of the effects of the factors, brain location, ctDCS montage, and state (ΔpretDCS, ΔtDCS, ΔpostDCS), on the canonical scores of O2Hb (Oxy, 3a and 3b) and the log10-transformed EEG bandpower (EEG, 3c and 3d). On each box in the box plot, the central mark indicates the median, and the bottom and top edges of the box indicate the 25th and 75th percentiles, respectively. The whiskers extend to the most extreme data points not considered outliers, and the outliers are plotted individually using the ‘+’ symbol.

Three-way ANOVA results on the effects of the factors, brain location, ctDCS montage, and state (Δ*pretDCS*, Δ*tDCS*, Δ*postDCS*), showed no significant effects on the NVC and the canonical scores of HHb at a 5% significance level. Boxplots of the NVC and the canonical scores of HHb are shown in Figure 4, where the central mark indicates the median, and the bottom and top edges of the box indicate the 25th and 75th percentiles, respectively. The whiskers extend to the most extreme data points not considered outliers, and the outliers are plotted individually using the ‘+’ symbol.

**Figure 4:**
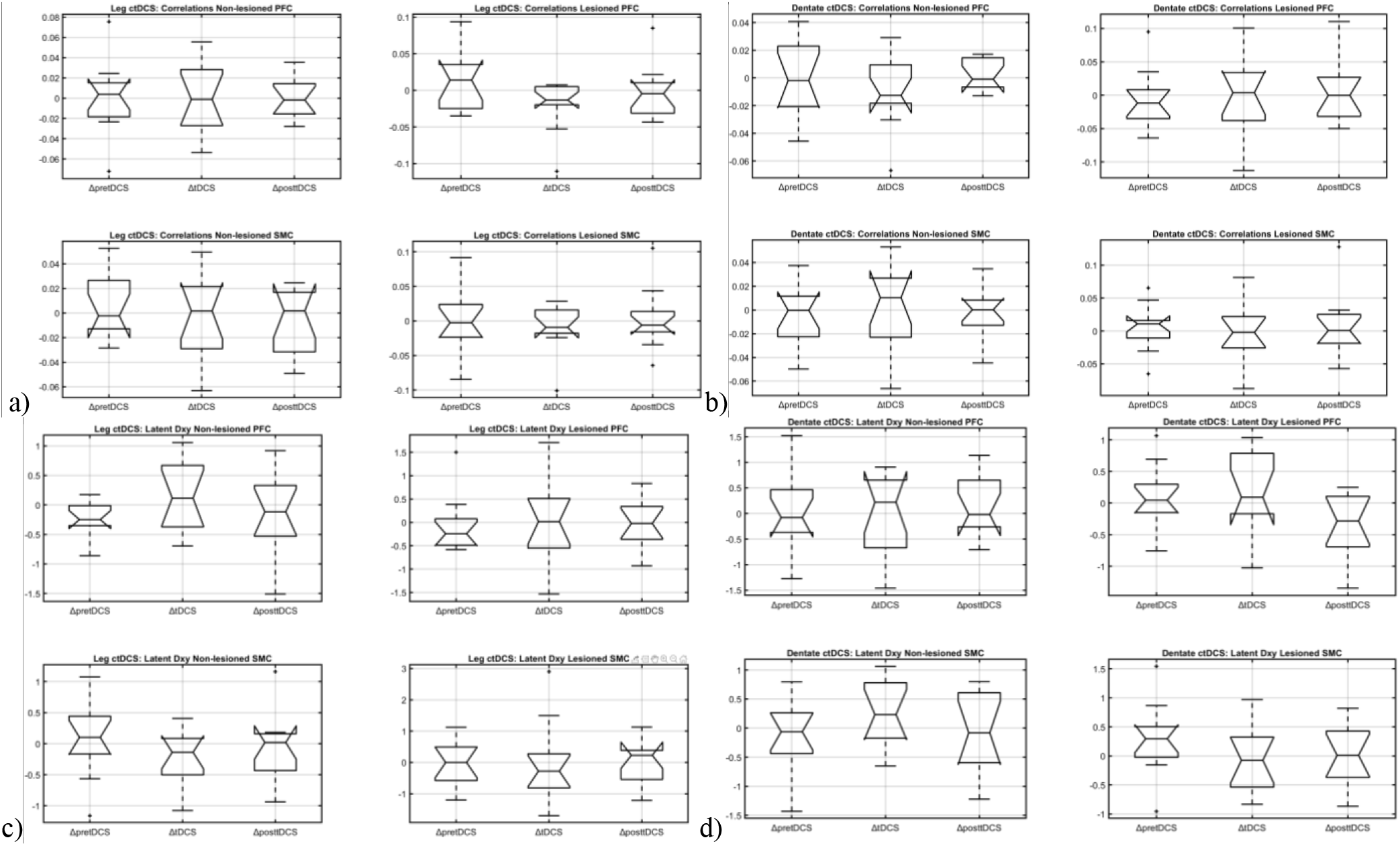
Boxplots of the effects of the factors, brain location, ctDCS montage, and state (ΔpretDCS, ΔtDCS, ΔpostDCS), on the canonical scores of NVC (Correlations, 4a and 4b) and HHb (Dxy, 4c and 4d) was found insignificant at a 5% significance level in a three-way ANOVA test. On each box in the box plot, the central mark indicates the median, and the bottom and top edges of the box indicate the 25th and 75th percentiles, respectively. The whiskers extend to the most extreme data points not considered outliers, and the outliers are plotted individually using the ‘+’ symbol.

### C. Multivariate general linear model (GLM)

In the case of mean lobular electric field strength, two principal components (PCs) accounted for greater than 98% variance (VAF) across the subjects. Figure 5a shows the hierarchical cluster tree (*dendrogram* plot) using the average Euclidean distance between 30 (without ‘Ipsi Interposed,’ ‘Contra Interposed,’ ‘Ipsi Fastigial,’ ‘Contra Fastigial,’ ‘Non-Cerebellar Brain’) pairs of mean lobular electric field strength across subjects that separated clusters for the dentate nucleus, and the cerebellar hemispheric lobules Crus I–Crus II along with the hemispheric lobules VIIb-IX. Figure 5b shows the first PC (PC1) of the mean lobular electric field strength (VAF= 97.5%) as the predictor and the latent variable of the O2Hb as the response variable (left panel) along with its standardized residuals (right panel). Figure 5c shows the first PC (PC1) of the mean lobular electric field strength (VAF= 97.5%) as the predictor and the latent variable of the log10-transformed EEG bandpower as the response variable (left panel) along with its standardized residuals (right panel). Here, both the standardized residual plots show a bivariate standard normal distribution with little evidence against the multivariate normality assumption.

**Figure 5:**
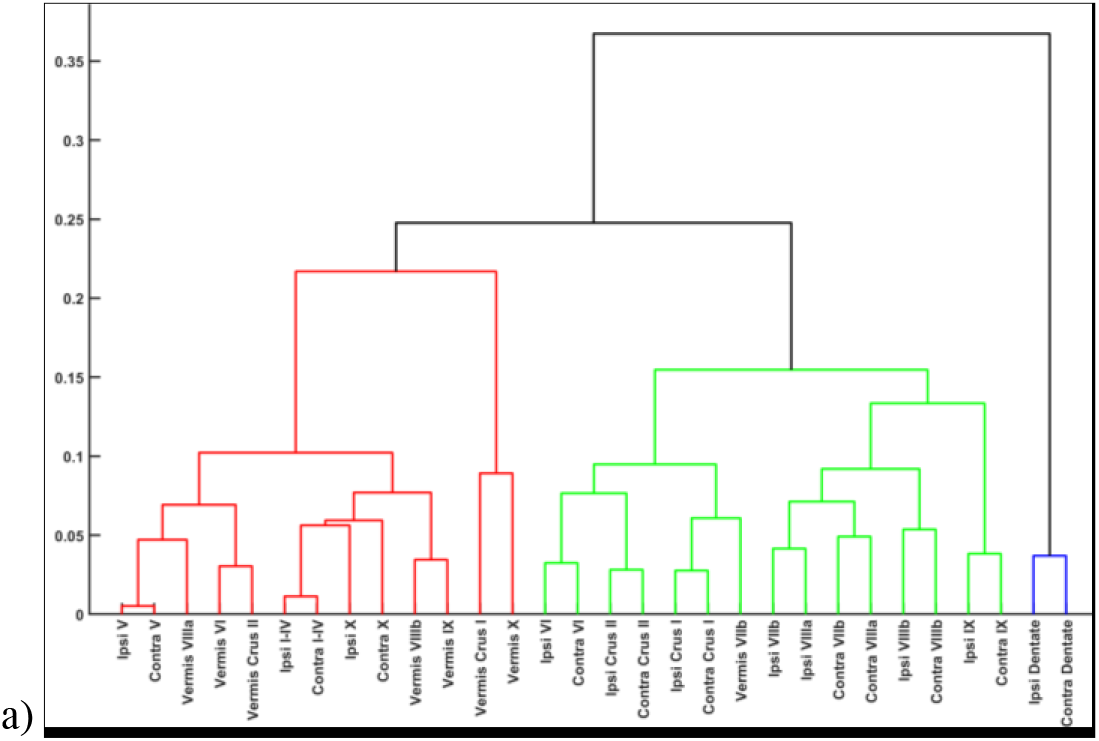
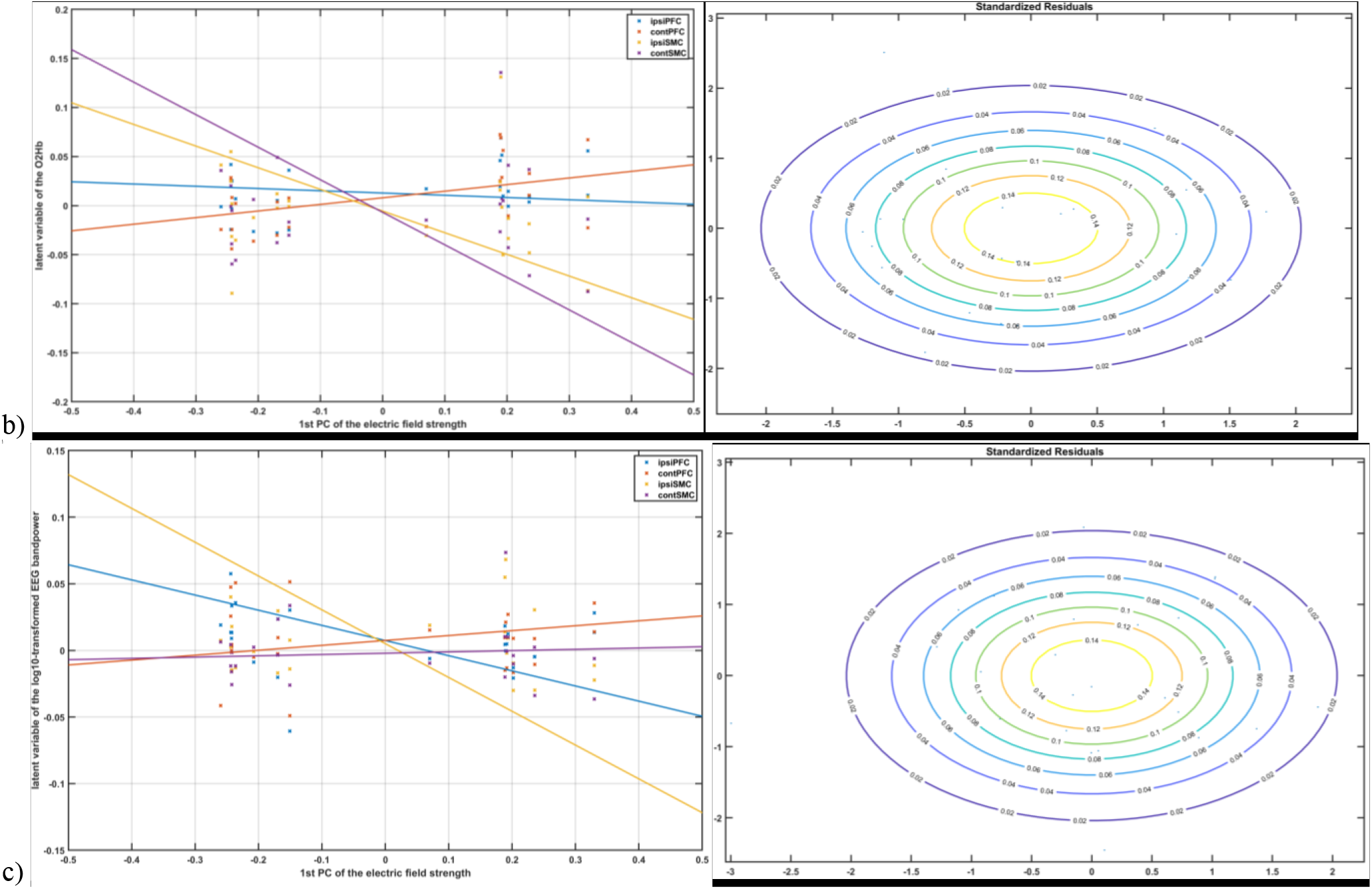
a) Dendrogram plot of mean lobular electric field strength across subjects and ctDCS montages where the x-axis corresponds to the leaf nodes of the tree, and the y-axis corresponds to the linkage Euclidean distance between the three main clusters (in red, green, blue), b) left panel shows the general linear model fit with the PC1 of the mean lobular electric field strength as the predictor and the latent variables of the O2Hb as the response. Here, ipsiPFC = 0.0128 + 0.0554 × PC1 − 0.0783 × PC2; contPFC = 0.0079 + 0.0758 × PC1 − 0.0085 × PC2; ipsiSMC = −0.0056 − 0.0199 × PC1 − 0.2011 × PC2; contSMC = −0.0068 + 0.0075 × PC1 − 0.3394 × PC2, c) left panel shows the general linear model fit with the PC1 of the mean lobular electric field strength as the predictor and the latent variables of the log10-transformed EEG bandpower as the response. Here, ipsiPFC = 0.0074 − 0.0228 × PC1 − 0.0910 × PC2; contPFC = 0.0075 − 0.0001 × PC1 + 0.0370 × PC2; ipsiSMC = 0.0051 + 0.0019 × PC1 − 0.2558 × PC2; contSMC = −0.0021 − 0.0023 × PC1 + 0.0120 × PC2.

## IV. Discussion

The computational neuroanatomy for motor control [78] suggested specific functions of different parts of the brain where the cerebellum builds internal models that predict the sensory outcome of motor commands and correct motor commands through internal feedback. Therefore, cerebellar role is important in an operant conditioning paradigm, e.g., to condition the usage of both the legs while interacting with the VBaT, thereby encouraging the hemiplegic stroke survivors for increased use of their paretic leg [79]. In contrast, primary motor (M1) and the premotor cortices are related to the implementation of the motor commands. This is achieved via cerebellar projections to M1 as well as premotor and other frontal regions [80]. Here, the premotor cortex has been implicated in the recombination and encoding of complex motor behaviors that are relevant for skill transfer. Prior works have shown that the interaction between the cerebellum and the M1 during motor skill learning is related to the cerebellar-M1 connectivity, estimated based on CBI, that decreases in the magnitude proportionally to the magnitude of motor skill learning [81]. This leads to dissociation in the processes of motor skill acquisition and retention where cerebellar tDCS has been shown to cause a rapid reduction of movement errors during skill acquisition. At the same time, M1 tDCS resulted in increased retention of the new skill [82]. In the current study, an interaction effect between the ctDCS montage and the state (pre-intervention, during ctDCS, post-intervention) was found on the change in the canonical scores of O2Hb (p=0.0788). Motor skill acquisition has been shown to decrease the activation of the prefrontal cortex [83], which was found in the canonical score of O2Hb at the non-lesioned PFC (and a small increase in SMC) in the novices following their first VBaT exposure before ctDCS intervention (i. e., Δ*pretDCS*) – see Figure 3a. Also, ctDCS with bipolar Exx7 – Exx8 montage for bilateral leg lobules VII-IX led to a decrease in the canonical scores of O2Hb at the lesioned SMC, which is postulated to be due to contra-lesional anode and CBI – see Figure 3a. Importantly, ctDCS intervention for bilateral leg lobules VII-IX led to an increase in the scores of O2Hb at the lesioned PFC and SMC, and a small decrease at the non-lesioned PFC and SMC, which is postulated to show increased use of the lesioned hemisphere during VBaT following ctDCS intervention. Moreover, significant effect of the state (p=0.0019) as well as the interaction between the ctDCS montage and the state (p=0.0153) was found on the changes in the canonical scores of log10-transformed EEG bandpower see Figure 3c, 3d. Hierarchical cluster analysis of the mean lobular electric field strength across all the subject headmodels showed three separate clusters for the dentate nucleus, and the cerebellar hemispheric lobules Crus I–Crus II along with the hemispheric lobules VIIb-IX for the two ctDCS montages– see Figure 5a. Here, the dentate nucleus cluster was found to be well separated from other two clusters which indicated a natural division across all the subject headmodels. The first PC (PC1) of the mean lobular electric field strength accounted for 97.5% of the variance (VAF) where PC1 was negatively correlated with the latent variables of the log10-transformed EEG bandpower at the contra-lesional PFC and SMC, and the ipsi-lesional PFC, while positively correlated with that for the ipsi-lesional SMC – see Figure 5c. For the latent variables of the O2Hb, the PC1 of the mean lobular electric field strength (VAF= 97.5%) was positively correlated with the latent variables of the O2Hb at the contra-lesional PFC and SMC, and the ipsi-lesional PFC, while negatively correlated with that for the ipsi-lesional SMC – see Figure 5b. These SMC effects are postulated to be due to ipsi-lesional cathode and contra-lesional anode (and CBI) where an increase in the bipolar ctDCS electric field strength led to an increase in the canonical score for log10-transformed EEG bandpower and a corresponding decrease in the canonical score for O2Hb (due to NVC) at the ipsi-lesional SMC while a decrease in the canonical score for log10-transformed EEG bandpower at the contra-lesional SMC (and a corresponding increase in the canonical score for O2Hb due to NVC). Here, a decrease in the canonical score for log10-transformed EEG bandpower can be related to a decrease in the SMC excitability [85]. In our knowledge, this is the first study reporting fNIRS-EEG based measure of brain activations at the PFC and SMC that can be putatively associated with the lobular electric field strength due to ctDCS therapy in the chronic stroke survivors.

We postulate that contra-lesional anodal cerebellar tDCS can facilitate “automatizing” of the post-stroke standing balance function where practice will lessen the cognitive demand, shifting peak fNIRS-EEG activation from PFC to the SMC. To investigate post-stroke standing balance function, CoP trajectories during cued weight shifts in different directions while interacting with the VBaT [19] elucidated ideomotor apraxia in our post-stroke subjects, which may contribute to patients’ overall day-to-day motor disability [69]. Here, VBaT tasks are goal-directed whole-body behavior using the CoP, where distinct neural circuits are responsible for the acquisition and performance of goal-directed behavior. Starting from the PFC, a cortical-dorsomedial striatal circuit is responsible for the acquisition of goal-directed actions while a cortical-ventral striatal circuit mediates the performance. Brain structures responsible for the goal-directed behavior are the medial prefrontal and orbitofrontal cortices, hippocampus, ventral, and dorsomedial striatum, whereas the sensorimotor cortices and dorsolateral striatum mediate the automatized/reflexive behavior in the experts. Here, the dorsomedial striatum (DMS) receives excitatory inputs from PFC whereas the dorsolateral striatum (DLS) primarily receives inputs from the sensorimotor and premotor cortices. This mapping from the prefrontal and sensorimotor cortices is important since DMS and DLS activity patterns have been shown [86] to be distinct during early training, but then they become similar following extended training leading to a more automatized (skilled) task performance. Therefore, it is postulated based on prior work [82] that anodal tDCS of the M1 can facilitate the retention of the learned sequence of movements for post-stroke standing balance function, which can be delivered following ctDCS facilitated motor learning. Since fNIRS-EEG imaging can provide a feedback of the shift in the peak activation from PFC to the SMC during ctDCS facilitated motor learning so fNIRS-EEG activation at the SMC can be maximally targeted with tDCS electric field strength based on the reciprocity principle [86], [87] after the subject reaches skilled performance. Direct fNIRS-EEG imaging of the cerebellum is not feasible, although fMRI has shown a decreased activation of the cerebellum with motor skill acquisition [83] so ctDCS can be turned off after the subject reached skilled performance. Our ongoing work is focused on the improvement of the fNIRS-EEG signal fidelity for monitoring the dynamical changes in brain activation associated with VBaT balance training. Also, we are investigating fNIRS-EEG based dynamic tDCS targeting of the moving fNIRS-EEG signal sources using current steering under reciprocity principle [86], [87]. This can allow staggered delivery of ctDCS for motor learning followed by M1 tDCS for skill retention based on the postulated shift in the peak activation from PFC to SMC in the responders. In the future, based on a large clinical study, fNIRS-EEG joint-imaging can also elucidate the inconsistency in ctDCS effects [40,41]. For example, ctDCS has shown promise in improving standing balance performance in small studies with fifteen patients with chronic stroke (>6 months post-stroke) [11] where exploration of optimal timing, dose, and the relation between qualitative parameters and clinical improvements are needed [11]. A recent study [20] showed that multiple sessions (three sessions of 20 min per week for two weeks) of simultaneous postural training with bilateral anodal ctDCS (not postural training or bilateral anodal ctDCS alone) was necessary to deliver therapeutic effects in older adults with high fall risk. A GLM analysis between the ctDCS electric field and latent variables for fNIRS-EEG coupling response may elucidate post-stroke functional brain connectivity that may correlate with white matter structural integrity in the corresponding cerebello-thalamo-cortical white matter tract [33]. Furthermore, fNIRS-EEG joint-imaging help to identify responders to ctDCS treatment based on the GLM analysis between the ctDCS electric field and the latent variables for EEG bandpower and fNIRS measures across subjects.

Limitations of this study include ‘one-size-fits-all’ ctDCS electrode montages; however, we performed MRI-guided computational modeling of the electric field using age-appropriate human brain MRI templates that were used as predictors in the multivariate GLM. We used subject matched age-groups for computational modeling to estimate electric field stimulation for the patients who had no lesion in the cerebellum. However, the differences in individual cerebellar anatomy and residual connectivity can play an essential role in the efficacy of the ctDCS [88], which may have resulted in the inter-subject variability in fNIRS-EEG results. Future clinical studies with a bigger sample size are required for unraveling the systematic effects of the optimized ctDCS for different cerebellar regions using a multivariate GLM. Here, the response variables in the multivariate GLM consisted of low-density (8-channel) fNIRS-EEG that provided limited coverage of the PFC and SMC. Due to a limited number of available fNIRS channels, we could not allocate short distance probes for the reduction of global interference of scalp-hemodynamics in our GLM to reduce fNIRS susceptibility to false positives [89]. Nevertheless, we postulate that the canonical scores for fNIRS O2Hb and HHb signal in the latent space are less sensitive to the systemic artifacts which needs to be evaluated in a future study. A limited 8-channel EEG prevented us from rejecting noisy channels during preprocessing in the EEGLAB toolbox [32], so we discarded all the VBaT task-related fNIRS-EEG data due to possible detrimental effects of co-modulated noise. We found good scalp coupling of the fNIRS optodes at the PFC based on PHOEBE approach that showed decreased activation in the non-lesioned PFC with increased VBaT at first exposure; however, the corresponding increase in the non-lesioned SMC activation was small which can be due to varying hair coverage and signal fidelity across subjects. Improvement of the sensitivity using silicon photomultipliers [90] can improve the fNIRS of the SMC. Nevertheless, low-density fNIRS-EEG allowed sensor placement on the head in a short amount of time that is important in a low-resource point-of-care setting for non-invasive electrical stimulation facilitated rehabilitation [91].

## Author Contributions

Neuroimaging in conjunction with non-invasive brain stimulation study conceptualization and design, A.D. (Anirban Dutta); Virtual reality-based balance training conceptualization and design, U.L. (Uttama Lahiri); Methodology, Z.R. (Zeynab Rezaee), D.S. (Dhaval Solanki), S.R. (Shashi Ranjan), A.D., and U.L.; Software development and application, Z.R., D.S., M.B. (Mahasweta Bhattacharya); Investigation, Z.R., D.S., and S.R.; Resources, A.D. (Anirban Dutta), U.L., and M.V.P.S. (MV Padma Srivastava); Data curation, Z.R., D.S., and S.R.; Writing—original draft preparation, Z.R., D.S., A.D., and U.L.; Writing—review and editing, A.D., U.L., and M.V.P.S.; Supervision, A.D., and U.L.; Project administration, A.D., U.L., and M.V.P.S; Funding acquisition, A.D., U.L., and M.V.P.S. All authors have read and agreed to the published version of the manuscript.

## Acknowledgment

Authors would like to acknowledge the technical support from Brandon Ruszala for the development of the CLOS pipeline and the clinical support from Surbhi Kaura for the MRIs and the stroke study at the All India Institute of Medical Sciences, New Delhi, India. Initial technology development and experimental validation of the VBaT platform was conducted by Deepesh Kumar during his doctoral research at the Indian Institute of Technology Gandhinagar, India. Authors also acknowledge the initial funding (2014−2017) by the Department of Science and Technology (DST), India and Institut National de Recherche en Informatique et en Automatique (Inria), France—https://team.inria.fr/nphys4nrehab/. This research is currently funded by the Indian Ministry of Human Resource Development (MHRD)′s Scheme for Promotion of Academic and Research Collaboration (SPARC), grant number 2018−2019/P721/SL, and Indian Department of Health Research, Project Code No. N1761.

